# NIMA-related kinase 1 (NEK1) regulates the localization and phosphorylation of α-Adducin (ADD1) and Myosin X (MYO10) during meiosis

**DOI:** 10.1101/110098

**Authors:** Miguel A. Brieño-Enríquez, Stefannie L. Moak, J. Kim Holloway, Paula E. Cohen

## Abstract

**Summary statement:** NEK1 kinase regulates the assembly and function of the meiosis I spindle by phosphorylating α-adducin (ADD1) and thereby facilitating its interaction with Myosin X (MYO10)

**Abstract:** NIMA-related kinase 1 (NEK1) is a serine/threonine and tyrosine kinase that is highly expressed in mammalian germ cells. Mutations in *Nek1* induce anemia, polycystic kidney and infertility. In this study we evaluated the role of NEK1 in meiotic spindle formation in both male and female gametes. Our results show that the lack of NEK1 provokes an abnormal organization of the meiosis I spindle characterized by elongated and/or multipolar spindles, and abnormal chromosome congression. The aberrant spindle structure is concomitant with the disruption in localization and protein levels of myosin X (MYO10) and α-adducin (ADD1), both of which are implicated in the regulation of spindle formation during mitosis. Interaction of ADD1 with MYO10 is dependent on phosphorylation, whereby phosphorylation of ADD1 enables its binding to MYO10 on mitotic spindles. Reduction in ADD1 protein in NEK1 mutant mice is associated with hyperphosphorylation of ADD1, thereby preventing the interaction with MYO10 during meiotic spindle formation. Our results reveal a novel regulatory role for NEK1 in the regulation of spindle architecture and function during meiosis.

## Introduction

Meiosis is a specialized cell division characterized by a single round of DNA replication followed by two rounds of chromosome segregation, resulting in the formation of haploid gametes for sexual reproduction. At the first male meiotic division (MI), homologous (maternal and paternal) chromosomes must segregate equally into two daughter cells, each of which then undergoes a mitosis-like second meiotic division (MII) in which sister chromatids separate into the haploid gametes. In mammals, meiosis in males results in four haploid gametes, while meiosis in female results in one haploid gamete per meiosis, the remaining genetic material being distributed between two polar bodies. Regardless of the sex of the individual, in order to achieve accurate segregation at both divisions, tension must be established on the meiotic spindles and this is achieved by the formation of crossovers between homologous chromosomes in meiosis I, and by cohesion between sister chromatids in both meiosis I and meiosis II (Gray and Cohen 2016). Cohesion is also necessary for sister kinetochore attachment to microtubules during meiotic spindle formation (Duro and Marston 2015).

Several families of kinases such as Polo kinases, Aurora kinases and NEK kinases have been implicated in the regulation of cell cycle events in both mitosis and meiosis (Fry et al. 2012, Meirelles et al. 2014). The founding member of the NEK family is the *Aspergillus nidulans* never-in-mitosis-gene-A (NIMA) protein (Oakley and Morris 1983). NIMA is a serine/threonine kinase involved in the initiation of mitosis and in promoting chromosome condensation. Genetic ablation of NIMA results in cell cycle arrest at G2, while overexpression of *NIMA* leads to premature entry into mitosis (Fry et al. 2012). In mammals, there are 11 orthologs of *NIMA* that comprise the NIMA-like (NEK) family of kinases. *Nek1* is unique in that it encodes both the serine/threonine kinase activity typical of the NEK family as well as tyrosine kinase activity that is not a feature of other NEKs (Letwin et al. 1992). NEK1 has been implicated in ciliogenesis (White and Quarmby 2008) and DNA damage response (Chen et al. 2011, Hilton et al. 2009, Liu et al. 2013, Patil et al. 2013). Additionally, NEK1 is highly expressed in mouse germ cells (Upadhya et al. 2000), where it appears to play essential roles in meiosis I and possibly also at later stages (Brieño-Enriquez et al. 2016, Holloway et al. 2011).

The *Kat2J* allele of *NEK1* is a spontaneous point mutation within the coding sequence of *Nek1* that results in a premature stop codon leading to a truncated protein lacking the both kinase domains (Upadhya et al. 2000). *Nek1^kat2j/kat2j^* mice display polycystic kidneys, dwarfism, anemia and male sterility (Janaswami et al. 1997, Upadhya et al. 2000). Our previous studies of meiosis in *Nek1^kat2j/kat2j^* mice showed that the loss of NEK1 activity induces aberrant retention of cohesin subunit Structural Maintenance of Chromosomes protein 3 (SMC3), at the end of meiotic prophase I (Holloway et al. 2011). More recently, we showed NEK1 regulates cohesin removal, in part, through regulation of wings apart-like homolog (WAPL) during meiotic prophase I (Brieño-Enriquez et al. 2016), as part of the so-called Prophase pathway for cohesin removal. This regulation of WAPL by NEK1 is mediated through WAPL interactions with the cohesin-associated protein, PDS5 homolog B (PDS5B), and protein phosphatase 1 gamma (PP1γ), both of which also interact with NEK1 (Brieño-Enriquez et al. 2016).

In the current study, and given the conserved function of NEK proteins in spindle assembly dynamics, we investigated the role of NEK1 downstream of prophase I events. Our results demonstrate that loss of NEK1 induces failure of meiotic spindle organization in both males and females, leading to elongated spindles, multipolar spindles, and abnormal chromosome congression. Concurrent with this, we observe altered levels and localization of two proteins known to regulate mitotic spindle dynamics: the unconventional myosin, myosin X (MYO10), and α-adducin (ADD1). Interaction of ADD1 with MYO10 is dependent on phosphorylation of ADD1 which allows for the interaction of these two proteins on mitotic spindles (Chan et al. 2014). Depletion of ADD1 or changes in its phosphorylation status results in abnormal mitotic spindles (Chan et al. 2014). Our studies demonstrate that loss of NEK1 kinase causes changes in MYO10 and ADD1 protein levels, and hyperphosphorylation of ADD1 caused by abnormal function of PP1 during meiotic spindle formation, independently of the role of NEK1 in prophase I events.

## Results

### Loss of NEK1 induces abnormal meiotic spindle formation and failed chromosome congression at the first meiotic division in males and females

We analyzed the role of NEK1 in spindle formation and chromosome orientation at MI in oocytes and spermatocytes from *Nek1*^*+/+*^ and *Nek1*^*kat2j/kat2j*^ mice. In male mice, we evaluated the shape, number of poles and misaligned chromosomes in both *Nek1*^*+/+*^ (n=100) and *Nek1*^*kat2j/kat2j*^ (n=100) spermatocytes. MI spindles from *Nek1*^*+/+*^ spermatocytes showed a bipolar structure with the chromosomes aligned at metaphase plate (Fig. 1A), while spindles from *Nek1*^*kat2j/kat2j*^ spermatocytes showed defects in the structure of the spindle as well as in chromosome congression. These defects included MI spindles without a pole, with only one pole, with misaligned chromosomes (Fig. 1B), and with multiple poles (Fig. 1C). A total of 55% of spindles from *Nek1*^*kat2j/kat2j*^ males were abnormal, while only 18% of the spindles from *Nek1*^*+/+*^ male spermatocytes showed abnormalities (Fig. 1D). The percentage of misaligned chromosomes in *Nek*^*+/+*^ spermatocytes was 18%, while in *Nek1*^*kat2j/kat2j*^ mice the percentage reached the 38% (Fig. 1E).

**Figure 1.**
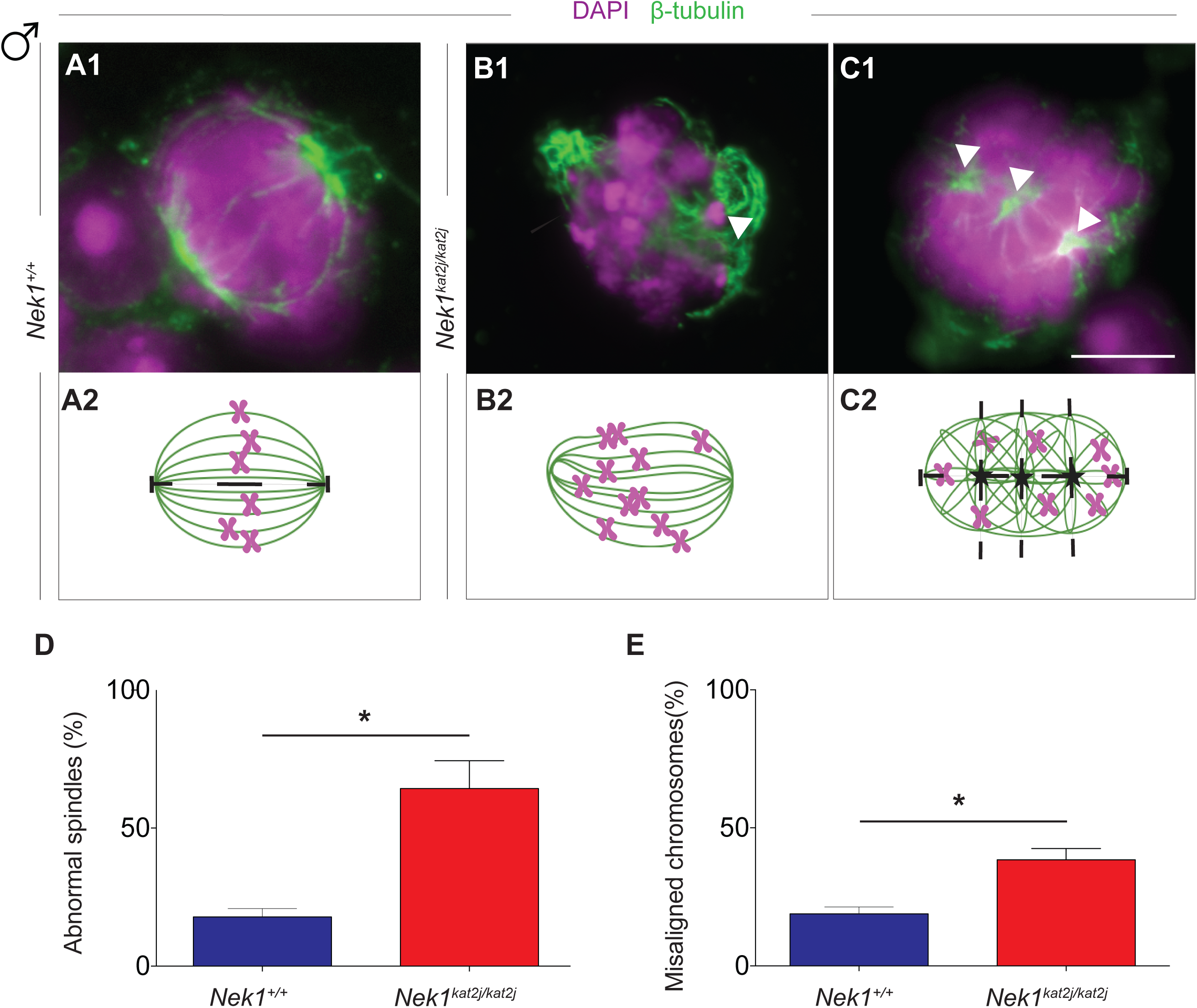
**Loss of NEK1 results in disrupted spindle morphology and chromosome
congression during meiosis I in spermatocytes.** (A-C) Immunofluorescence (IF) against β-tubulin (green) and DNA staining with DAPI (magenta) showing meiotic spindle morphology (panels A1-C1), with graphic representations below (panels A2-C2). (A) *Nek1*^*+/+*^ spermatocyte shows normal spindle morphology and chromosome congression at the midplate of the spindle; (B, C) Examples of aberrant spindle morphology in spermatocytes from *Nek1*^*kat2j/kat2j*^ mice showing monopolar spindles with mislocalized chromosomes resulting from failed chromosome congression (arrowheads) in B and multipolar spindles in C (arrowheads). (D, E) Quantitation of abnormal spindles and misaligned chromosomes in spermatocytes from *Nek1*^*+/+*^ and *Nek1*^*kat2j/kat2j*^ mice (n=100 cells counted for each). Values are percentages ± Standard deviation.* Indicates statistically significant differences (Unpaired t-test p < 0.05).

The analysis of MI spindles in oocytes from *Nek1*^*+/+*^ females showed a bipolar structure of the spindle and appropriate congression of the chromosomes at the metaphase plate (Fig. 2A). However, MI spindles from *Nek1*^*kat2j/kat2j*^ females showed a distortion of the structure of the spindle characterized by the presence of multiple spindles, mini spindles and misaligned chromosomes (Fig 2B, 2C). We evaluated the percentage of abnormal spindles in both wildtype oocytes (n=100) and mutant oocytes (n=100). In *Nek1*^*kat2j/kat2j*^ female spindles, 82% of the MI spindles were abnormal, while only 22% of the *Nek1*^*+/+*^ oocytes showed any abnormality (Fig. 2D). Similarly, 88% of MI spindles from *Nek1*^*kat2j/kat2j*^ oocytes showed at least one misaligned chromosome, compared to only 23% in oocytes from *Nek1*^*+/+*^ mice (Fig. 2E).

**Figure 2.**
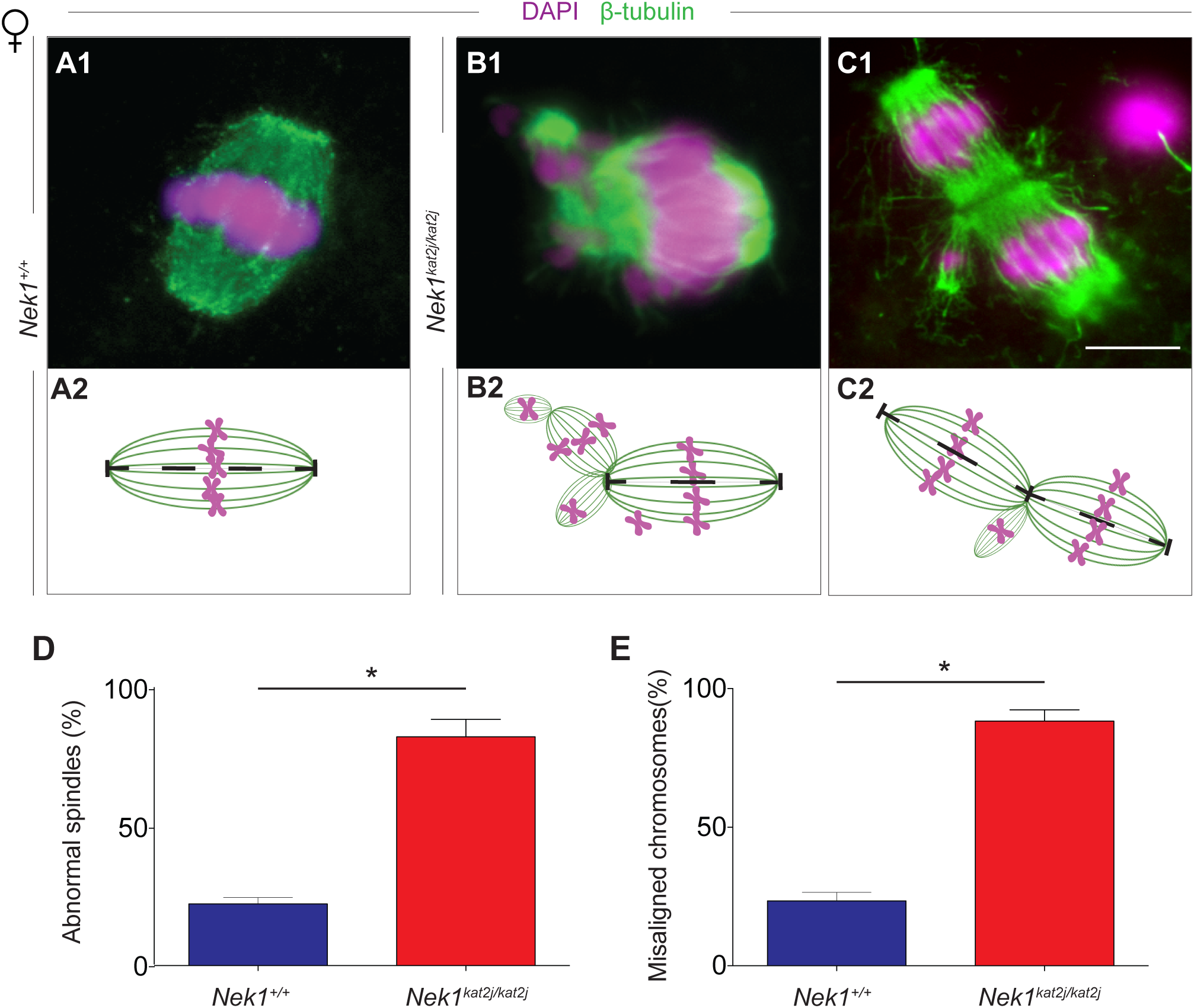
**Loss of NEK1 results in disrupted spindle morphology and chromosome
congression during meiosis I in oocytes.** (A-C) Immunofluorescence (IF) against β-tubulin (green) and DNA staining with DAPI (magenta) showing meiotic spindle morphology (panels A1-C1), with graphic representations below (panels A2-C2). (A) *Nek1*^*+/+*^ oocyte shows normal spindle morphology and chromosome congression at the midplate of the spindle; (B, C) Examples of aberrant spindle morphology in oocytes from *Nek1*^*kat2j/kat2j*^ mice showing multipolar spindles with or without errors in chromosome congression (arrowheads indicate three extra spindles formed in the oocyte in B and two additional spindles in C). (D, E) Quantitation of abnormal spindles and misaligned chromosomes in oocytes from *Nek1*^*+/+*^ and *Nek1*^*kat2j/kat2j*^ mice (n=100 cells counted for each). Values are percentages ± Standard deviation. * Indicates statistically significant differences (Unpaired t-test p < 0.05).

### *Nek1*^*ka2tj/kat2j*^ spermatocytes and oocytes have an abnormal spindle length and width

The appropriate length and width of the spindle are critical for establishing tension during segregation, and thus both parameters were evaluated them in male and female spindles. We measured the distance between poles in MI spermatocyte spindles and we observed a significant increase of the length of *Nek1*^*kat2j/kat2j*^ males (27.9 µm ± 7.0 s.d.) compared to that of *Nek1*^*+/+*^ males (23.1 μm ± 2.0 s.d.; Unpaired t test p=0.0020; Fig. 3A). Increases in MI spindle length in male spermatocytes were also accompanied with increases in the width in MI spindles from *Nek1*^*kat2j/kat2j*^ male mice (26.7 μm, ± 6.4 s.d.) compared to that of *Nek1*^*+/+*^ male mice (22.9 μm ± 2.1 s.d; Unpaired t test p=0.0097; Fig. 3B). We also analyzed the length and width of the female spindles. *Nek1*^*kat2j/kat2j*^ oocytes showed an increase in the length of the spindle in MI oocytes (31.6μm ± 7.3 s.d.) compared to that found in *Nek1*^*+/+*^ females (26.0 μm ± 2.3 s.d.; Unpaired t test p=0.0007; Fig. 3C). The lack of NEK1 also resulted in altered width of the MI spindles in female meiosis. *Nek1*^*kat2j/kat2j*^ oocytes showed a decrease in the width of the spindle (15.1 μm ± 4.0 s.d.) compared to the *Nek1*^*+/+*^ oocyte spindles (17.0 μm ± 1.7 s.d.; Unpaired t test p=0.039; Fig. 3D). Taken together our results show that NEK1 is required for the correct spindle formation and chromosome alignment during meiosis in both male and female gametes.

**Figure3.**
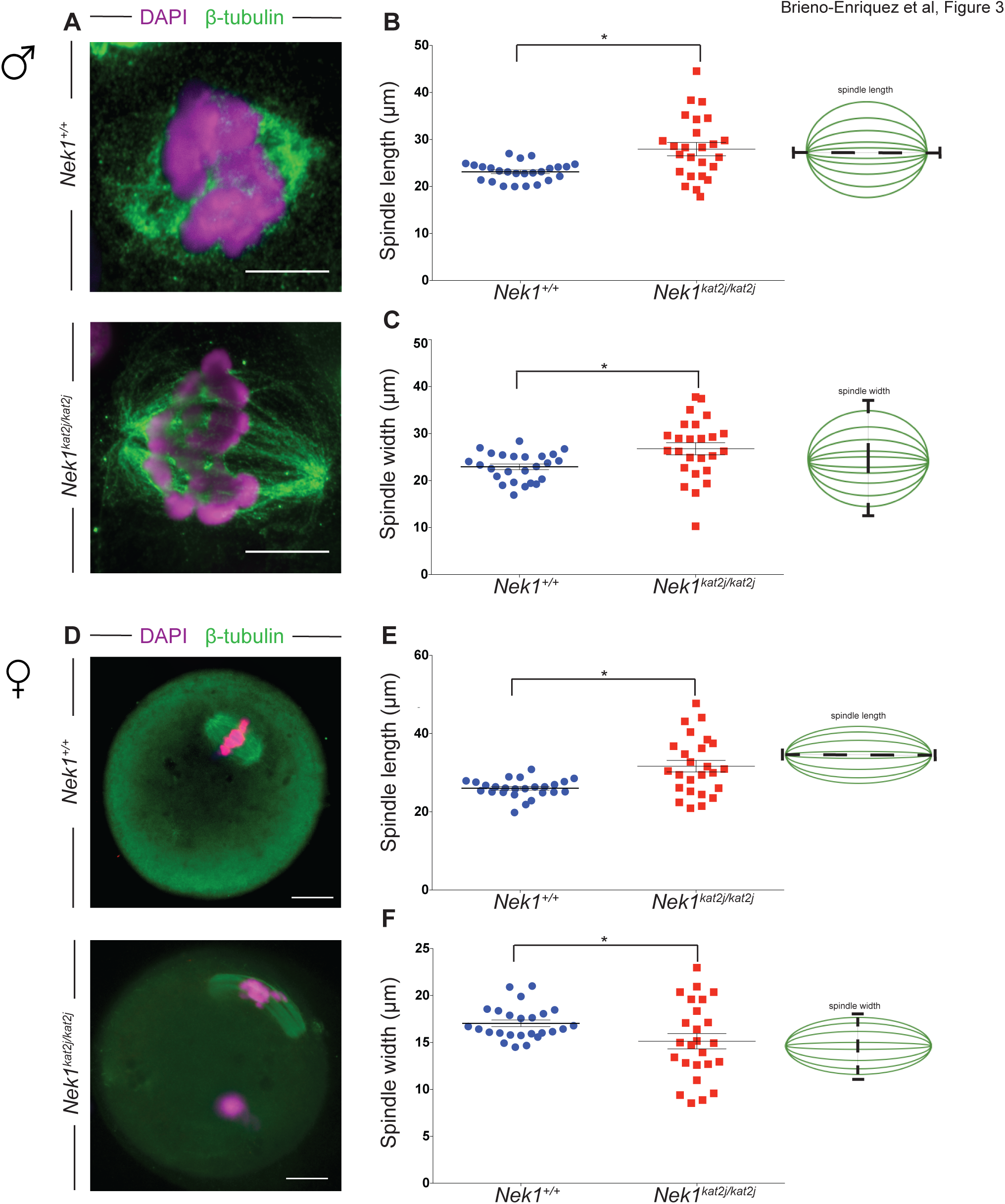
**Lack of NEK1 results in abnormal meiosis I spindle morphology in spermatocytes and oocytes**. (A) Immunofluorescence (IF) against β-tubulin (green) and DNA staining with DAPI (magenta) showing a wildtype (top) and *Nek1*^*kat2j/kat2j*^ spermatocytes (bottom), along with measurements of spindle length (B) and width (C). Black bars are means (μm) ± Standard deviation. (D) Immunofluorescence (IF) against β-tubulin (green) and DNA staining with DAPI (magenta) showing a wildtype (top) and *Nek1*^*kat2j/kat2j*^ (bottom) oocytes, along with measurements of spindle length (B) and width (C). Black bars are means (μm) ± Standard deviation. * Indicates statistically significant differences (Unpaired t-test p < 0.05).

### Lack of NEK1 results in deregulation of myosin X (MYO10) and α-adducin (ADD1)

Given the disrupted spindle apparatus during meiosis I, we investigated possible protein targets of NEK1 to explain this phenotype. In a previous study, we had performed Mass spectrometry (MS) screening using testis protein extracts from *Nek1*^*+/+*^ and *Nek1*^*kat2j/kat2j*^ mouse (Brieño-Enriquez et al. 2016). Review of these data using GO terms for centrosome and centromeric proteins revealed no significant differences in these proteins levels in *Nek1*^*kat2j/kat2j*^ mouse extracts compared to wildtype mice (Fig. S1). Several proteins associated with spindle formation and regulation were identified and quantified, however no differences between *Nek1*^*+/+*^ and *Nek1*^*kat2j/kat2j*^ mouse were observed for tubulins, actins and SAC proteins (Fig. S2, S3, S4).

Previous reports showed that MYO10 has a direct function in meiotic spindle formation (Sandquist et al. 2016, Weber et al. 2004, Woolner et al. 2008). In contrast to the tubulins, actins, and other spindle proteins that did not appear to be altered in the absence of NEK1, we observed that the levels of the myosins, MYO5, MYO7A, MYO10 and MYO15, were significantly increased in testis protein extracts from *Nek1* mutant mice compared to wildtype littermates (Table S1). By contrast, the myosins, MYO1B, MYO1C, MYO1D, MYO7A and MYO9B were unaltered. The function of MYO10 in spindle formation and integrity is related to its binding partner α-adducin (ADD1)(Chan et al. 2014). Our MS results revealed that ADD1 showed lower protein levels in *Nek1*^*kat2j/kat2j*^ mouse compared to wildtype mouse (Table S1). The changes in protein levels observed by MS indicates that the loss of NEK1 results in an imbalance in the ratio MYO10/ADD1, and this could be affecting the proper spindle formation and function.

### *Nek1*^*kat2j/kat2j*^ spermatocytes and oocytes show aberrant localization of myosin X (MYO10) on meiotic spindles

We evaluated the localization of MYO10 on MI spindles from male spermatocytes and female oocytes. Immunofluorescence (IF) analysis in *Nek1*^*+/+*^ spindles revealed that MYO10 localizes along the β-tubulin fibers specifically in the region where tubulin interacts with the chromosome (the kinetochores) in mouse spermatocytes (Fig. 4A). We analyzed the number of MYO10 foci per nucleus in the coronal plane (Fig. 4B) and found an increase in the number of foci in the *Nek1*^*kat2j/kat2j*^ mice (75.4 ± 6.7 s.d.) compared to *Nek1*^*+/+*^ animals (66.3 ± 5.5 s.d.; Unpaired t test p=0.0001). Analysis of transverse sections of male spindles revealed that there is a subtle accumulation of MYO10 at the spindle pole in *Nek1*^*+/+*^ male mice (Fig. 4A), and this was disrupted in *Nek1*^*kat2j/kat2j*^ mice (Fig. 4B). In the transverse plane, we observed an increase in the MYO10 foci number in *Nek1*^*kat2j/kat2j*^ mice (74.7 ± 5.4 s.d.) compared to the Nek1^+/+^ mice (61.7 ± 4.4 s.d.; Unpaired t test p=0.0001). The increase in MYO10 focus counts was supported by our mass spectrometry analysis showing increased MYO10 protein in testis extracts from *Nek1*^*kat2j/kat2j*^ males compared to *Nek1*^*+/+*^ mice (Table S1).

**Figure 4.**
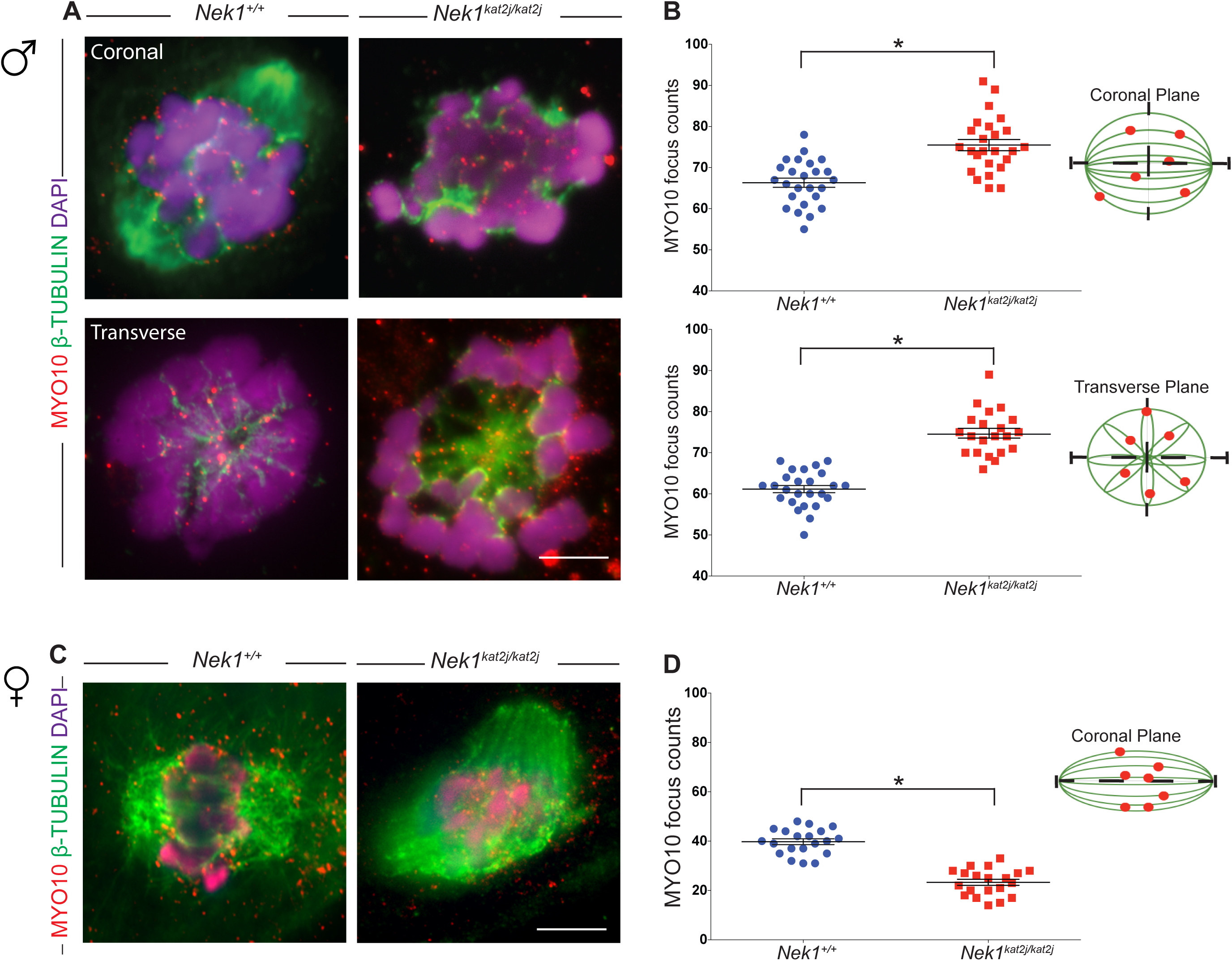
***Nek1*^*kat2j/kat2j*^ spermatocytes and oocytes show aberrant localization of myosin X (MYO10) on meiosis I spindles.** (A) IF against MYO10 (red), β-tubulin (green) and DAPI (magenta) on spermatocytes from wildtype (left) and *Nek1*^*kat2j/kat2j*^ mice (right) in coronal (top) and transverse (bottom) planes. Arrowheads indicate examples of MYO10 foci. Quantitation of MYO10 focus counts per nucleus are given in panel (B). (C) IF against MYO10 (red), β-tubulin (green) and DAPI (magenta) in oocytes of both wildtype and mutant mice. Arrowheads indicate examples of MYO10 foci. Quantitation of MYO10 focus counts per nucleus are given in panel (D). All values are focus counts per spindle and black bars denote means ± Standard deviation. * Indicates statistically significant differences (Unpaired t test p < 0.05).

We also evaluated the MYO10 foci in *Nek1*^*+/+*^ oocytes (Fig. 4C), MYO10 localizes on tubulin fibers as foci during MI (39.7 ± 5.1 s.d.). However, in oocytes obtained from *Nek1*^*kat2j/kat2j*^ mouse, the number of MYO10 foci that colocalize with β-tubulin is significantly reduced (23.3 ± 5.4 s.d.; Unpaired T test p=0.0001)(Fig. 4D). Taken together, our results indicate that loss of NEK1 leads to mislocalization of MYO10 along meiotic spindles in both male and females germ cells.

### Localization of ADD1 on spermatocytes and oocytes is disrupted in absence of NEK1

Our previous MS results showed that ADD1 proteins levels are reduced in testis extracts from *Nek1*^*kat2/kat2j*^ mouse compared to wildtype littermates (Table S1)(Brieño-Enriquez et al. 2016). Thus, we evaluated the localization and protein levels of ADD1 in both male and female spindles during meiosis I. ADD1 localizes both to the poles of the spermatocyte spindles (Fig. 5A, asterisk) and with β-tubulin fibers of the spindle (Fig. 5A, arrow heads). We evaluated the ADD1 focus number associated with the spindle structure in the coronal and transverse planes. MI spindles from *Nek1*^*kat2j/kat2j*^ males showed a decreased ADD1 focus number (25.7 ± 7.8 s.d.) compared to MI spindles from *Nek1*^*+/+*^ males (40.3 ± 3.2 s.d.), along with mislocalization of ADD1 on the spindle poles on the coronal plane (Unpaired t test p=0.001)(Figure 5B). Analysis of the transverse plane of spindles from *Nek1*^*kat2j/kat2j*^ males showed a decreased focus number (32.6 ± 4.6 s.d.) compared to that of spindles from *Nek1*^*+/+*^ male (22.3 ± 5.5 s.d.; Unpaired t test p=0.0001). Reduced ADD1 focus numbers were supported by reduced ADD1 protein levels analyzed by MS in testis extracts from NEK1 mutant mice and wildtype littermates (Table S1). In oocytes from *Nek1*^*+/+*^ females, ADD1 also localized to the spindle poles as two large foci on each pole (3.6 ± 0.9 s.d.)(Fig. 5C, 5D). This localization pattern was disrupted in *Nek1* mutant females, where we observed multiple foci on the poles (6.08 ± 1.4 s.d.; Unpaired t test p=0.0002) indicating the mislocalization of the protein could be result as a pole fragmentation like in somatic cells (Chan, et al., 2014).

**Figure 5.**
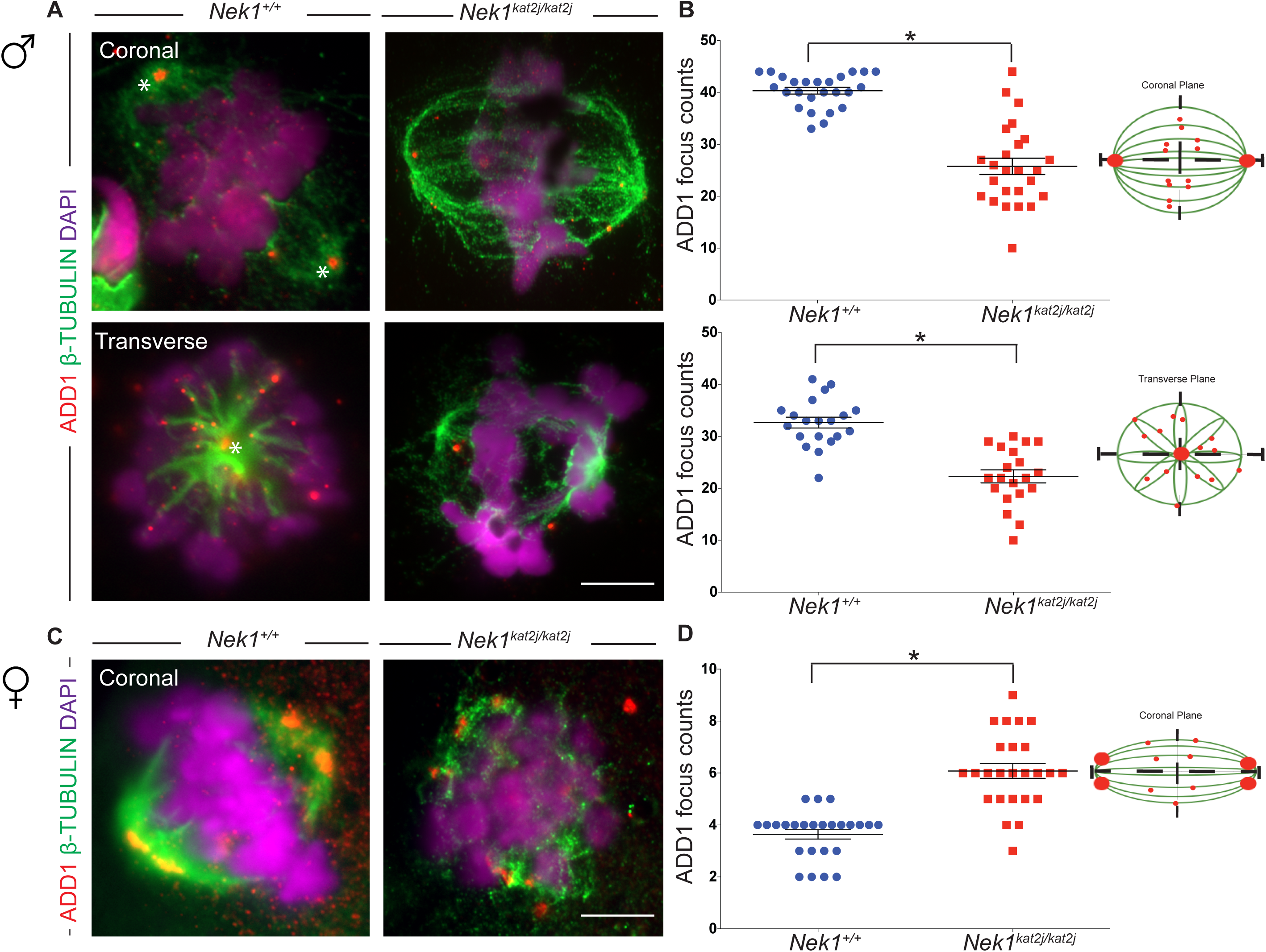
***Nek1*^*kat2j/kat2j*^ spermatocytes and oocytes show aberrant localization of Adducin 1 (ADD1) on meiosis I spindles.** (A) IF against ADD1 (red), β-tubulin (green) and DAPI (magenta) on spermatocytes from wildtype (left) and *Nek1*^*kat2j/kat2j*^ mice (right) in coronal (top) and transverse (bottom) planes. Arrowheads indicate examples of ADD1 foci. Quantitation of ADD1 focus counts per nucleus are given in panel (B). (C) IF against ADD1 (red), β-tubulin (green) and DAPI (magenta) in oocytes of both wildtype and mutant mice. Arrowheads indicate examples of ADD1 foci. Quantitation of ADD1 focus counts per nucleus are given in panel (D). All values are focus counts per spindle and black bars denote means ± Standard deviation. * Indicates statistically significant differences (Unpaired t test p < 0.05).

### Loss of NEK1 activity induces abnormal phosphorylation of ADD1

During mitotic spindle formation mammalian cell lines, ADD1 binds to the motor domain of MYO10 on the spindle. ADD1 and MYO10 interaction on mitotic spindles is negatively regulation by phosphorylation. Phosphorylation of ADD1 induces the loss of the interaction between both proteins and therefore the complex unloads from the spindle (Chan et al. 2014). Phosphorylation of ADD1 results in abnormal meiotic spindle morphology (elongation, multipolar spindles and aberrant chromosome alignment) and loss of its interaction with MYO10 (Chan et al. 2014). We evaluated the proteins levels of ADD1 in testis lysates and isolated oocytes. In both male and female germ cells, we observed a double band in *Nek1*^*+/+*^ lysates (both bands were included for quantitation), however this double band is reduced or not present in protein lysates from *Nek1*^*-/-*^ testes and oocytes. Our WB results revealed a decrease in the protein levels in testis from mutant mice compared to wildtype testis and oocyte lysates (Fig. 6A; Unpaired t test p=0.0031). Decreased levels of ADD1 and the loss of bands in WB analysis of *Nek1*^*kat2j/kat2j*^ mouse compared to wildtype littermates suggested that the lack of NEK1 activity affects ADD1 phosphorylation, leading to a reduced interaction with MYO10. To test this hypothesis, we screened our previous phosphoproteomics data obtained from *Nek1* mutant animals, specifically focusing on changes in ADD1. Loss of NEK1 results in a significant increase in the phosphorylation of ADD1 (Fig. 6B)(Brieño-Enriquez et al. 2016). Specifically, loss of NEK1 results in hyperphosphorylation of ADD1 on serine 465 (S465) (Fig. 6C). In oocytes the WB analysis revealed a decrease in the protein levels in *Nek1*^*kat2j/kat2j*^ oocytes compared to *Nek1*^*+/+*^ oocytes (Fig. 6D; Unpaired t test p=0.0003). To evaluate the changes in the ADD1 protein levels at various times of oocyte culture, we observed ADD1 protein levels at 6 hours after initial culture. Interestingly, in *Nek1*^*+/+*^ oocytes we observed the loss of the heavier band indicating a change in ADD1 phosphorylation. However in the *Nek1*^*kat2j/kat2j*^ oocytes we observe that both bands almost disappear indicating that there is a change in the phosphorylation in the mutants but also degradation of the protein in the mutant oocytes.

**Figure 6.**
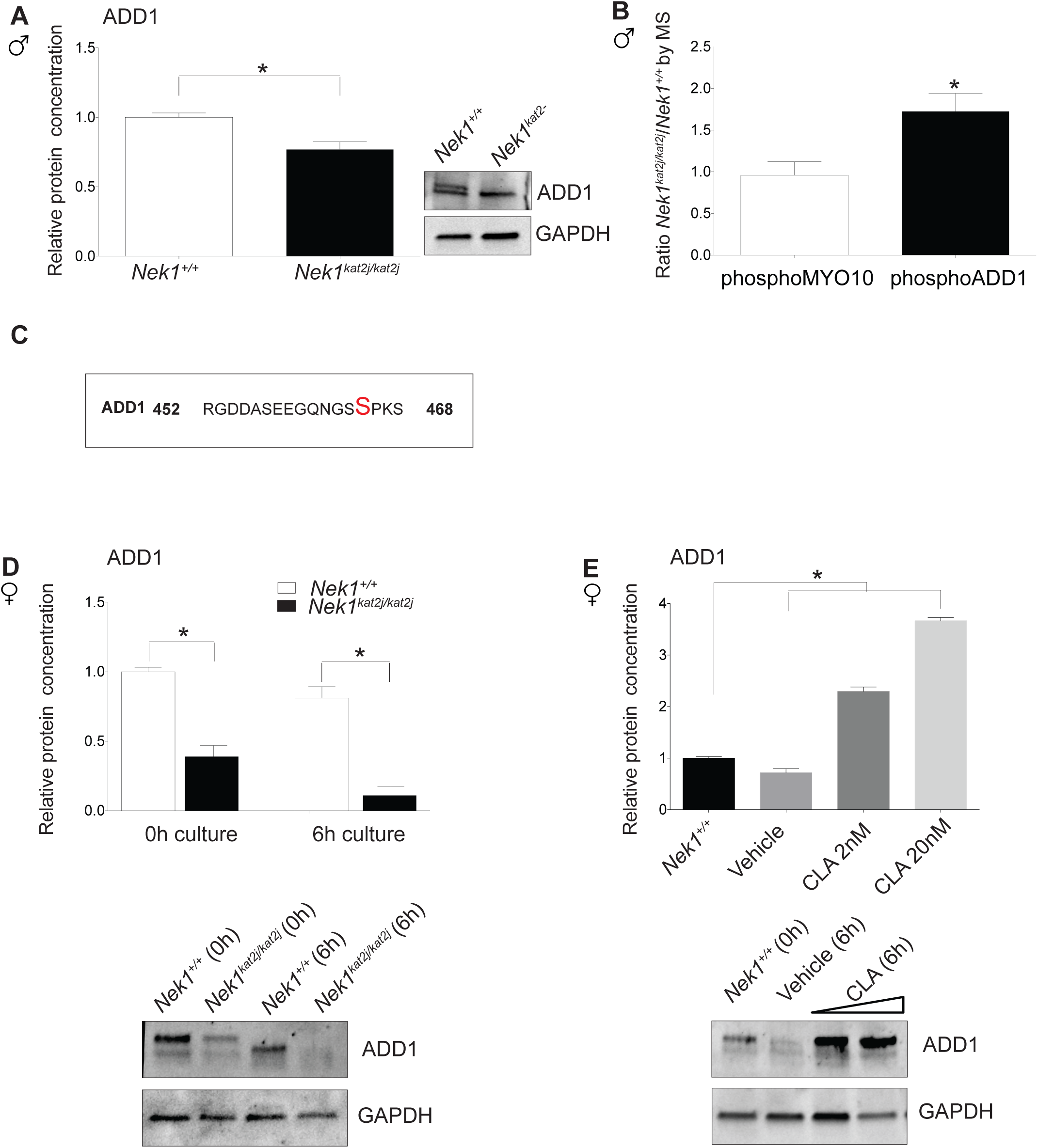
**Loss of NEK1 during meiosis is associated with hyper-phosphorylation and reduced ADD1 protein in a PP1γ-dependent manner.** (A) Quantitation of ADD1 protein levels in *Nek1*^+/+^ testis extracts relative to GAPDH control. (B) ADD1 and MYO10 protein levels, expressed as a ratio of *Nek1*^*kat2j/kat2j*^ / *Nek1*^*+/+*^, as determined by mass spectrometry. (C) ADD1 phosphopeptide sequence determined by mass spectrometry to be higher in *Nek1*^*kat2j/kat2j*^ testis extracts. (D) Quantitation by western blotting of ADD1 protein levels in oocytes at 0h and after 6h of culture. (E) Quantification of ADD1 protein levels in wildtype oocytes cultured for 6h in vehicle (ethanol) and the PP1 inhibitor Calyculin A (CLA) (at doses of 2 and 20 nM). All values are means ± Standard deviation. * Indicate statistically significant differences (Unpaired t test p < 0.05 and one-way ANOVA followed by Dunnett’s multiple comparisons test, p < 0.05, for cultures with inhibitor)

We previously reported the interaction of PP1γ with NEK1 and its function regulating the prophase pathway during meiosis (Brieño-Enriquez et al. 2016). To test if the hyper-phosphorylation of ADD1 depends on PP1 activity, we cultured *Nek1*^*+/+*^ oocytes for 6h in the presence of the PP1 inhibitor, Calyculin (Fig.6E). The WB analysis of these cultures showed that the inhibition of PP1 induces a statistically significant increase in the ADD1 protein levels, predominantly the heavier band, suggesting that the inhibition of PP1 maintains the phosphorylation status of ADD1. By contrast, loss of NEK1 loss did no affect MYO10 phosphorylation status, suggesting that the effects on MYO10 protein may be indirect, through ADD1. Thus, these results suggest that the lack of NEK1 results in increased ADD1 phosphorylation that is associated with a disruption in ADD1-MYO10 interactions.

## Discussion

In eukaryotes, the structure orchestrating chromosome alignment and segregation during cell division is the microtubular spindle (Bennabi et al. 2016). To ensure correct spindle dynamics, the spindle assembly checkpoint (SAC) becomes activated in situations where tension is not appropriately established between the chromosomes and the spindle poles, and this will block the progression of the cell cycle to prevent aberrant segregation. In addition to the canonical SAC, other levels of control have been described, including MYO10 (Sandquist et al. 2016, Weber et al. 2004, Woolner et al. 2008). In oocytes and frog embryos, disruption of MYO10 negatively affects nuclear anchoring and meiotic spindle formation (Weber et al. 2004), while depletion of MYO10 in frog epithelial cells induces abnormal spindle movements, spindle elongation, multi-spindle and pole fragmentation (Kwon et al. 2015, Woolner et al. 2008). MYO10 has an amino-terminal globular head domain that harbors acting biding and ATPase activities, giving to the protein the capacity to bind F-actin. The tail of MYO10 has the MyTH4 and FERM domains that give to the protein the ability to interact with microtubules (Hirano et al. 2011, Weber et al. 2004). Accordingly, recent studies have demonstrated that loss of *Myo10* results in spindle dysfunction, while overexpression of only the MyTH4 domain alone affects the structure and shape of the spindle (Sandquist et al. 2016).

In the current study, we present evidence to suggest that loss of NEK1 induces increases in MYO10 protein levels, and that this in turn may result in spindle defects. Our interpretation that overexpression of *Myo10* leads to spindle defects is in line with the suggestion that over-expression of the MyTH4 domain of MYO10 results in competition with the wildtype protein, leading to the displacement of the latter from the spindle (Hirano et al. 2011). Furthermore, it has been suggested that over-expression of only the MyTH4 may disrupt the functional link between F-actin, microtubules and MYO10. In our data (Table S1), we observed an increase in the MYO10 protein levels without changes in F-actin or microtubules that could be inducing an imbalance in the protein levels provoking a similar phenotype to the overexpression of the MyTH4 domain.

ADD1 is an actin-biding protein that is important for membrane stabilization (Anong et al. 2009) and for cell-cell adhesion (Abdi and Bennett 2008, Franco et al. 2016). There are 3 isoforms of ADD with similar domain structures that are formed by NH_2_-terminal head domain, a neck domain and a C-terminal domain. The C-terminal domain is characterized by it high contain of myristoylated alanine-rich C kinase substrate (MARCKS), this MARCKS related domain is necessary for its interaction with F-actin, spectrin, and calmodulin (Hirano et al. 2011, Kaiser et al. 1989, Matsuoka et al. 1996, Scaramuzzino and Morrow 1993). Previous studies in somatic cells showed that loss of ADD1 leads to a failure in mitotic spindle formation characterized by distortion, elongation, multipolar spindles and abnormal chromosome alignment (Chan et al. 2014). Here, we show that loss of NEK1 leads to a reduction in ADD1 protein levels, abnormal ADD1 distribution on the meiotic spindle, and consequent disruption of spindle formation and integrity. These phenotypes observed in meiotic cells are highly reminiscent of the phenotypes observed in somatic cells lacking *Add1* (Chan et al. 2014).

ADD1 and MYO10 interact by the MyTH4 domain of the MYO10, and this interaction is critical for the correct spindle formation and function (Chan et al. 2014, Hirano et al. 2011, Weber et al. 2004, Woolner et al. 2008). Interaction of ADD1 to MYO10 is dependent on phosphorylation; phosphorylation of ADD1 by cyclin-dependent kinase 1 (CDK1) enables ADD1 to bind MYO10 at mitotic spindle (Chan et al. 2014). Depletion of ADD1 or changes in phosphorylation status results in abnormal mitotic spindles (elongation, multipolar spindles and aberrant chromosome alignment) (Chan et al. 2014). In the absence of NEK1 we observe an increase in the phosphorylation of S465. Such modification could induce a premature loss of the ADD1-MYO10 interaction similar to the phenotypes observed in somatic cells (Chan et al. 2014). However, it is unclear what kinase is phosphorylating ADD1 during meiosis, since the increased phosphorylation in the absence of NEK1 suggests the up-regulation of a kinase that is itself regulated by NEK1 (either directly or indirectly). Clear candidates could be CDKs, however, our MS results did not show any significant change in the profile of CDK in the absence of NEK1, and specifically no differences in CDK1 indicating that increases in phosphorylation of ADD1 does not depend of CDK1 pathway. Alternatively, the increased phosphorylation of ADD1 could suggest the existence of a phosphatase that is inactivated by the loss of NEK1, allowing the hyperphosphorylation of protein. We previously described that in *Nek1* mutants, there is a down regulation and abnormal phosphorylation in whole testis lysate of PP1γ, and our results of oocyte cultures in the presence of a PP1 inhibitor suggest that the increase in ADD1 phosphorylation could be mediated by a similar mechanism.

In summary, we propose that the loss of NEK1 activity induces abnormal spindle formation through a mechanism mediated by the imbalance of MYO10 and ADD1. The decreases in ADD1 and it abnormal localization on the meiotic spindle is mediated by an increase its phosphorylation that leads to loss of interaction of MYO10 with ADD1. Taken together our results show the importance of NEK1 in the control of spindle formation during male and female meiosis.

## Material and Methods

### Mice and genotyping

All mouse studies were conducted with the prior approval of the Cornell Institutional Animal Care and Use Committee (Protocol 2010-0054). The *Nek1*^*kat2j/kat2j*^ line was obtained originally from Jackson Laboratory (Bar Harbor, Maine), and has been established in our mouse colony for more than nine years. Genotyping of this mouse line was performed following the protocol described elsewhere (Holloway et al. 2011). For the purposes of these studies, male mice were 8 weeks old and female where used at 24-28 days old. Homozygous mutant animals of (*Nek1*^*kat2J/kat2J*^) were compared with wild type (*Nek1*^*+/+*^) littermates on a C57Bl/6J background.

### Male and female spindle preparation

Male spindle analysis was performed in testis from 8 week old mice following the protocol described elsewhere (Kotaja et al. 2004, Page et al. 1998). Briefly, following euthanasia, the testes were removed, decapsulated and placed in PBS. Using fine forceps, tubules were separated and analysed under a stereomicroscope. Differential light absorption of the tubules creates different zones that can be defined by different cell populations (Kotaja et al. 2004). The differentent zones are consequence of the paracrine regualtion by Sertoli cells, spermatogenesis proceeds in synchronized waves along the seminiferus tubules, and every given cross-section contains only certain cell types, that produces a diferent light absorbation creating zones (Kotaja et al. 2004). These zones are the weak spot that correspond to the spermatogenic stages XII-I, the strong spot (stages II-VI), the dark zone (stages VII-VIII) and the pale zone (stage IX-XI) We selected the zone corresponding to the spermatogenetic development XI and XII that are rich in meiotic cells during both meiosis I and meiosis II. Tubules were placed in fresh PBS and these required zones were excised and placed on poly-lysine-coated slides (P4981, Thermo Fisher). Tubule sections were fixed for 10 minutes (1% paraformaldehyde and 0.15% Triton-X, pH 9.2) and then squashed under a coverslip. Slides with the squashed tubules were put into liquid nitrogen for 20 seconds and thereafter the coverslip removed and washed with PBS-0.4% Kodak Photo flo (Kodak) (3 times for 5 minutes each). Slides were kept at −80°C or stained immediately.

Female spindle analysis was performed with oocytes from 24-28 day old mice following our previously published protocol (Sun and Cohen 2013). Briefly, after dissection, ovaries were collected in collection medium (Waymouths’s medium (11220035, Gibco), 10% Fetal bovine serum (26140079, Gibco), 1% penicillin-streptomycin (P4333-100ML, Sigma-Aldrich) and 0.1% sodium pyruvate (11360070, Gibco)). Oocytes were released from ovarian follicles using 30-gauge needles, and oocytes with germinal vesicles placed in EmbryoMax KSOM medium (MR-121-D, EMD Millipore). Oocytes were cultured for 6hr in EmbryoMax KSOM medium drops at 37°C, 5% CO2. After incubation, oocytes were placed in fibrinogen-thrombin clot, incubated for 60 seconds at 37°C, and rinsed with 1XPBS containing 2% Triton-X (BP151-500, Fisher Scientific) for 3 min. Oocytes were fixed for 30 minutes at 37°C. Fixation was followed by a 15 minute wash in 0.1% NGS (5 ml 10X PBS, 50 μL goat serum and 50 ml MIlli-Q water). Oocytes were stored at 4°C or stained immediately.

### Spindle immunofluorescence staining

Male and female fixed spindles slides were washed in PBS containing 0.4%Kodak Photo flo (Kodak) for 5 min, followed by 0.1% PBS-Triton X and blocked in 1XPBS-antibody dilution buffer (ADB) before being incubated over night at 4°C with the primary antibody. Primary antibodies used were: rabbit anti-ADD1 (GTX101600, Genetex. Dilution 1:100), rabbit anti-MYO10 (24565-1-AP, Proteintech. Dilution 1:100) and rabbit anti-β-tubulin (T8328, Sigma. Dilution 1:500). After overnight incubation, slides were washed to remove the unbound antibodies and incubate for 2 hours at 37°C with Alexafluor™ secondary antibodies (Molecular Probes Eugene OR, USA). Slides were washed and mounted with Prolong Gold antifade (Molecular Probes). Image acquisition was performed using a Zeiss Imager Z1 microscope under 20X, 40X or 63X magnifying objectives, at room temperature. Images were processed using ZEN 2 (Carl Zeiss).

### Mass Spectrometry

Mass spectrometry was performed in the Cornell University Proteomics and Mass Spectrometry facility. 2D LC-MS/MS raw data files were acquired using Orbitrap Elite (Thermo Scientific). We performed a database search using Mascot searching against the SwissProt mouse database from Uniprot website (http://www.uniprot.org) using Mascot software version 2.3.02 (Matrix Science, UK). The default Mascot search settings were as follows: one missed cleavage site by trypsin allowed with fixed MMTS modification of cysteine, fixed four-plex iTRAQ modifications on Lys and N-terminal amines and variable modifications of methionine oxidation, deamidation of Asn and Gln residues, and 4-plex iTRAQ on Tyr for iTRAQ 4-plex analysis. One or two-missed cleavage site by trypsin allowed with fixed carboxamidomethyl modification of cysteine, fixed six-plex TMT modifications on Lys and N-terminal amines and variable modifications of methionine oxidation, deamidation of Asn and Gln residues, and 6-plex TMT on Tyr for TMT 6-plex analysis. The quantitative protein ratios were weighted and normalized by the median ratio with outlier removal set automatic in Mascot for each set of experiments. Only those proteins with ratios equivalents to two-fold increase or two-fold reduction were considered significant. To obtain the specific post-traslational modification of the peptides we perfumed TiO2 enrichment followed by the proteomics analysis as was described above.

### Western blotting

Whole testis protein and oocyte protein were extracted by sonication in RIPA buffer. Samples were boiled for 5 min in sample buffer, electrophoresed on SDS-polyacrylamide gels (8%) and transferred to nitrocellulose membranes. Primary antibody incubation was performed overnight at 4°C at 1:1000 dilution (antibodies are the same used in spindle staining). Incubation with secondary antibodies was performed for two hours at room temperature (secondary HRP conjugated antibodies were obtained from Pierce, Life Technologies). Signal-detection was carried out using the superSignal substrate (Thermo Scientific). Loading control was performed using GAPDH-HPR (PA1-987-HRP, from ThermoFisher). Images were captured with BIO RAD Image Lab 5.1and analyzed by ImageJ version 1.49v (http://rsbweb.nih.gov/ij).

## Acknowledgements

We are grateful to Peter Borst for assistance in animal care and genotyping. We thank Dr. Sheng Zhang from Cornell Proteomics and Mass Spectrometry Facility for his advice in development of mass spectrometry experiments. This project is funded by grants from the NICHD to J. K. H (5R00HD065870), and from NIGMS and March of Dimes to P. E. C. (1R01GM097263, MOD2006-844).

### Competing interests

The authors declare no competing interest

### Author contribution

M.A.B-E, J.K.H and H P.E.C designed experiments. M.A.B-E and S.L.M carried out experiments. M.A.B-E, S.L.M, and P.E.C analyzed and interpreted data. M.A.B-E and P.E.C wrote the manuscript.

---

**Figure S1.**
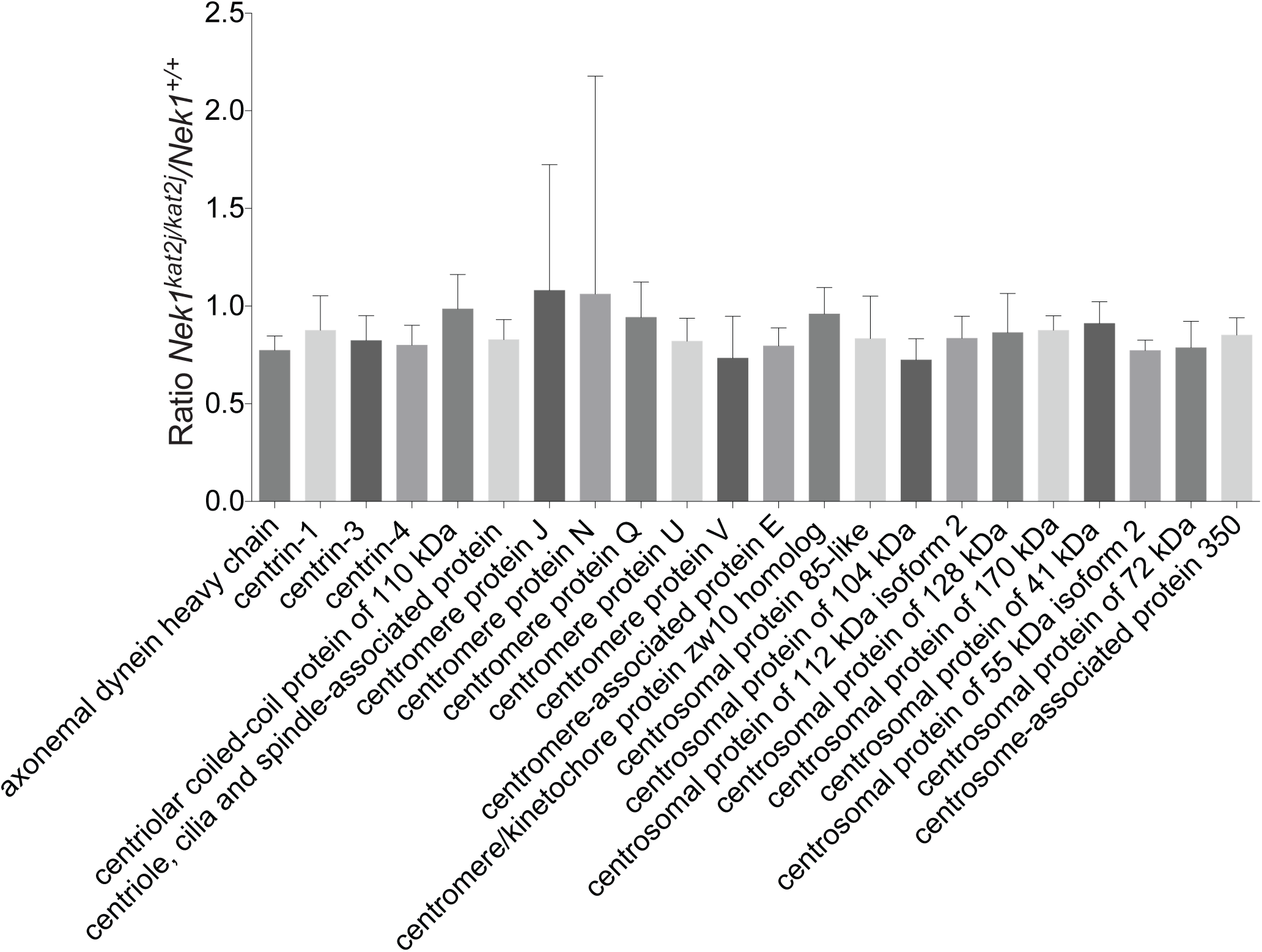
**Analysis of centromere and centrosomal protein ratio *Nek1^kat2j/kat2j^/Nek1^+/+^* by mass spectrometry in whole testis lysates.**

**Figure S2.**
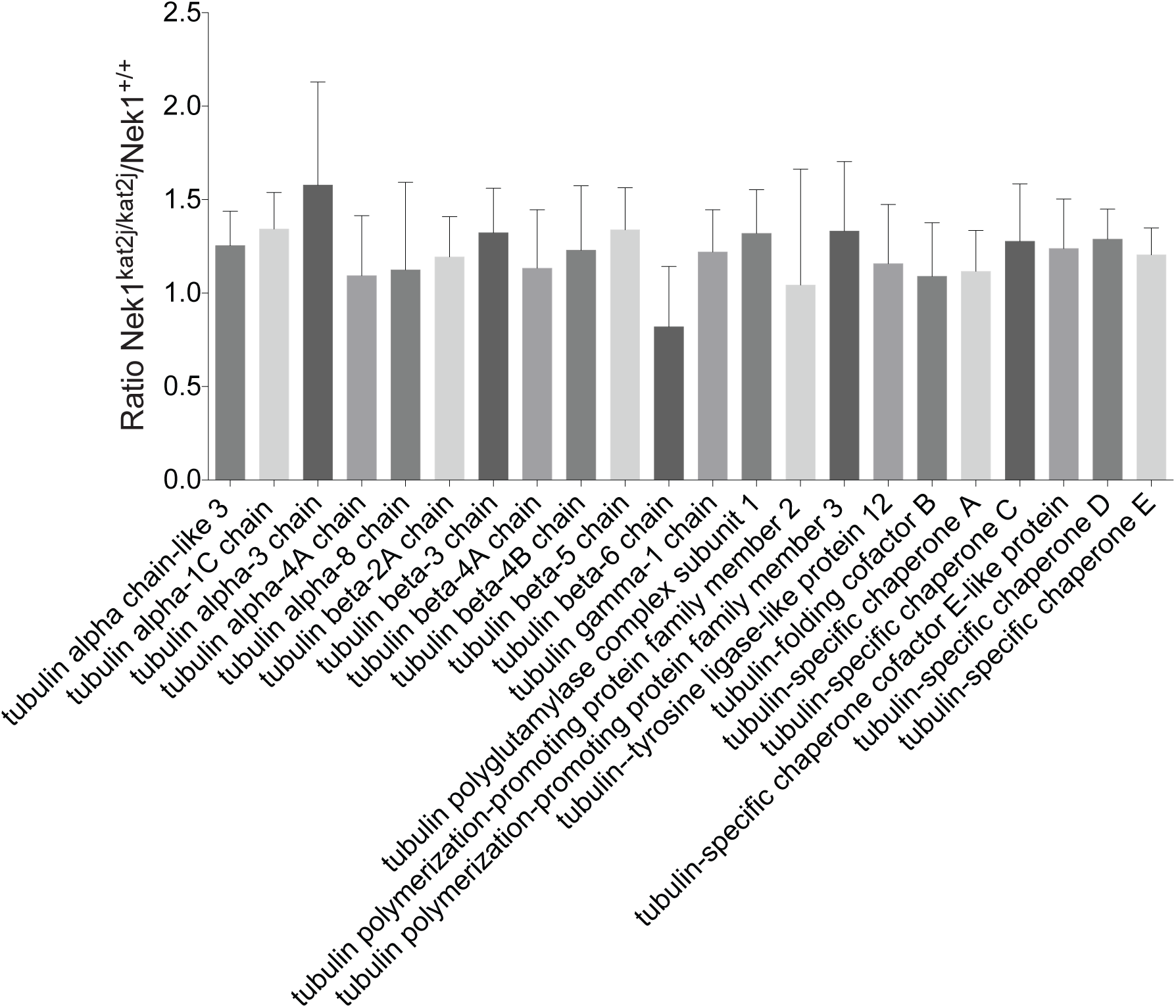
**Analysis of Tubulins protein ratio *Nek1^kat2j/kat2j^ /Nek1^+/+^* by mass spectrometry in whole testis lysates.**

**Figure S3.**
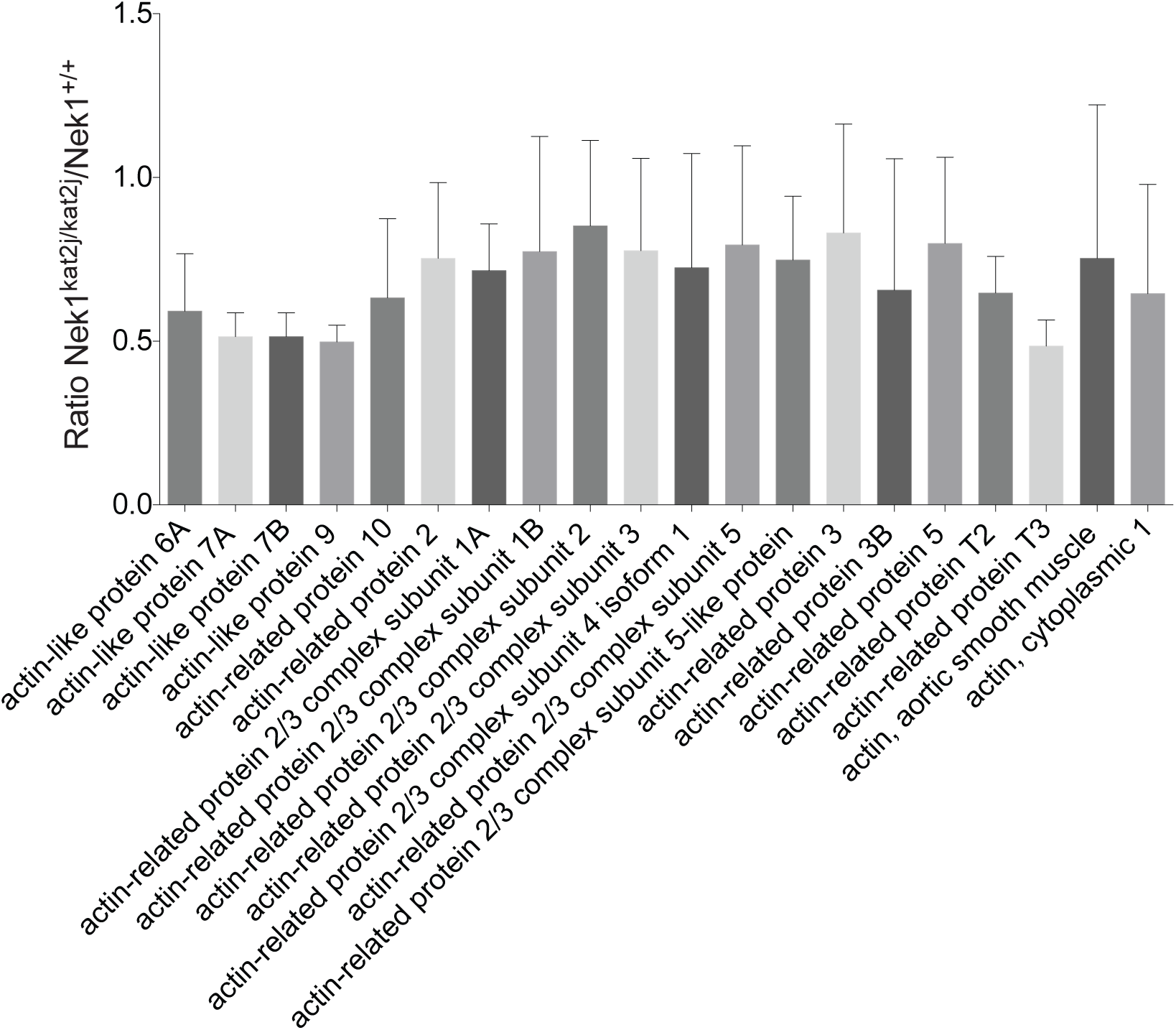
**Analysis of Actins protein ratio *Nek1^kat2j/kat2j^ /Nek1^+/+^* by mass spectrometry in whole testis lysates.**

**Figure S4.**
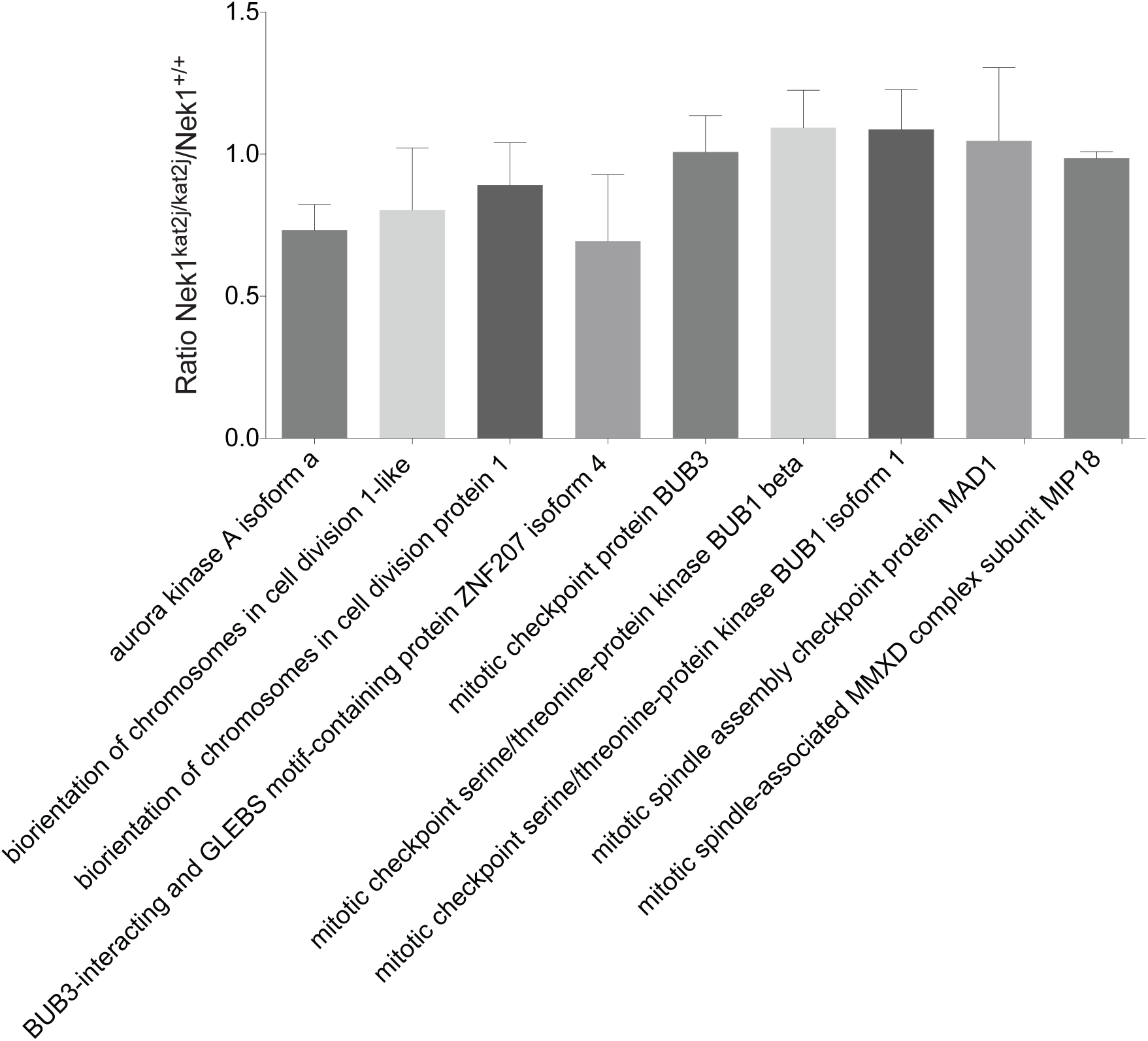
**Analysis of Spindle assembly checkpoint protein ratio *Nek1^kat2j/kat2j^ /Nek1^+/+^* by mass spectrometry in whole testis lysates.**

**Table S1.** **Myosin proteins quantitation by mass spectrometry**

|  | **Mean** | **SD** |
| --- | --- | --- |
| myosin light chain 1/3, skeletal muscle isoform isoform 1f | 1.234777778 | 0.583235325 |
| myosin light chain 4 | 0.916222222 | 0.189561849 |
| myosin light chain 6B | 0.945222222 | 0.221693921 |
| myosin light chain kinase 3 isoform 1 | 1.277333333 | 0.550780129 |
| myosin light chain kinase, smooth muscle | 1.284555556 | 0.30677317 |
| myosin light chain, regulatory B-like | 1.005888889 | 0.193254654 |
| myosin light polypeptide 6 | 1.195333333 | 0.296617178 |
| myosin regulatory light polypeptide 9 | 1.086 | 0.287876276 |
| myosin-10 | 1.061222222 | 0.102613081 |
| myosin-11 isoform 2 | 1.110444444 | 0.408137266 |
| myosin-14 isoform 3 | 1.212333333 | 0.480961017 |
| myosin-7 | 1.131222222 | 0.285432031 |
| myosin-9 | 1.218111111 | 0.627280528 |
| myotubularin-related protein 1 | 1.04275 | 0.056646145 |
| myotubularin-related protein 5 isoform 1 | 1.023222222 | 0.159993576 |
| myotubularin-related protein 6 | 1.006888889 | 0.126838918 |
| myotubularin-related protein 9 | 0.910888889 | 0.144002122 |
| PREDICTED: myosin phosphatase Rho-interacting protein isoform X8 | 0.897777778 | 0.176093567 |
| PREDICTED: unconventional myosin-XVIIIa isoform X17 | 0.999888889 | 0.066242819 |
| unconventional myosin-Id | 1.199888889 | 0.353706957 |
| unconventional myosin-IXb isoform 3 | 1.149666667 | 0.447813856 |
| unconventional myosin-Va | 1.24 | 0.353951974 |
| unconventional myosin-Vb | 1.636333333 | 0.528680669 |
| unconventional myosin-VI | 1.457222222 | 0.338891862 |
| unconventional myosin-VIIa isoform 2 | 1.535 | 0.16826913 |
| unconventional myosin-X | 1.810222222 | 0.38393319 |
| unconventional myosin-XV isoform 1 | 1.914 | 0.490053313 |
| alpha-adducin isoform 1 | 0.605111111 | 0.052415276 |

